# Distinct domain requirements for EAP45 in HIV budding, late endosomal recruitment, and cytokinesis

**DOI:** 10.1101/2020.05.23.112607

**Authors:** Bo Meng, Pedro P. Vallejo Ramirez, Katharina M. Scherer, Ezra Bruggeman, Julia C. Kenyon, Clemens F. Kaminski, Andrew M. Lever

## Abstract

The scission of lipid membranes is a common biological process, often mediated by ESCRT complexes in concert with VPS4 assembling around the separation point. The functions of the ESCRT-I and ESCRT-III complexes are well established in certain of these cellular processes; however, the role of ESCRT-II remains contentious. Here, we devised a SNAP-tag fluorescent labelling strategy to understand the domain requirements of EAP45, the main component of ESCRT-II, in HIV egress, late endosome recruitment, and cytokinesis. We used TIRF microscopy to measure the spatial co-occurrence of the HIV structural polyprotein Gag with full length EAP45 in both fixed and live cells. Gag colocalises with the full length EAP45 comparably to ALIX, but this is lost on deletion of the EAP45 N terminus. Our findings reveal the H0 domain of the EAP45 protein is essential for linking to ESCRT-I during HIV budding and in anchoring at the late endosomal membrane, however in cytokinesis it is the Glue domain that is critical.

## Introduction

ESCRTs (endosomal sorting complexes required for transport) are protein complexes involved in a variety of cellular functions, from the scission of the intercellular bridge during cytokinesis, to the formation of multi-vesicular bodies (MVB) and the extracellular budding of a number of viruses including human immunodeficiency virus (HIV). ESCRT activities involve the topological remodelling of membranous structures. The core of the ESCRT machinery comprises ESCRT-I (TSG101, VPS28, VPS37, and MVB12/UBAP1), -II (EAP20, EAP30, and EAP45) and -III (CHMP1–7) acting in a sequentially coordinated manner during membrane scission (1). The ESCRT machinery also has two other associated complexes, ESCRT-0 which precedes early cargo recruitment, and VPS4 AAA ATPase which universally acts at a late stage for membrane scission (2).

Many viruses take advantage of the properties of the ESCRTs during their life cycle. In the case of HIV, a small number of Gag polyproteins form oligomers in the cytosol, possibly with the genomic RNA, before they are trafficked to the plasma membrane where further Gag molecules nucleate and membrane deformation occurs (3–5). Two late domains within the p6 domain of Gag, PTAP and YPXL, interact with the ESCRT-I protein TSG101 and ESCRT-associated protein ALIX, respectively, which then recruit ESCRT-III for the final scission to occur (3). Our understanding of the recruitment of the ESCRTs during Gag accumulation at the membrane has benefitted from biochemical analyses of budding events (6–9), as well as from advanced fluorescence microscopy techniques which can examine the temporal dynamics of these events (10–13). Interestingly, TSG101 and ALIX show different dynamics once engaged with Gag; TSG101 co-occurs with Gag as Gag multimerises (14), whereas ALIX is only recruited when Gag puncta reach maximum intensity (13). ESCRT-III components are recruited to the budding site and remain there for 3-5 mins (10) with recent data indicating recurrent transient recruitment of CHMP4B and VPS4 until successful cleavage occurs (5). Due to the transitory nature of these recruitments, the temporal co-occurrence between various components of ESCRT with Gag is very low, with only ∼1-4% of the total Gag at the plasma membrane detectably colocalised with an ESCRT component, as measured by direct stochastic optical reconstruction microscopy (dSTORM)(15).

Extensive work has been carried out in elucidating the function of ESCRT-I and ESCRT-III in HIV budding and in other cellular processes (16), however the role of ESCRT-II remains unclear. ESCRT-II comprises one copy of EAP30 and EAP45 (Fig. 1A) and two copies of EAP20 forming a Y shaped structure (17). Structural studies in yeast show ESCRT-I interacts with ESCRT-II via the C-terminal domain (CTD) of VPS28 and the N terminus of the VPS36 (EAP45 in metazoans) Glue and H0 linker domains(1, 17, 18) (Fig. 1A). A homologous region in mammalian EAP45 was also shown to interact with TSG101 by yeast two-hybrid screening (19). The remaining domains of EAP45 (WH1 and WH2) and EAP30 form the core structure of the ESCRT-II complex with EAP20 (17, 20), which is responsible for recruiting ESCRT-III (17), though an alternative link is also documented (21). Although ESCRT-II’s role as the canonical bridge in linking to ESCRT-I in MVB formation (22, 23) and cytokinesis is established (24), the same role has been controversial in HIV budding. An earlier siRNA knockdown study of EAP20 suggested ESCRT-II is dispensable for HIV budding in 293T cells (19), however, our recent study using CRISPR/ EAP45 knockout (KO) HAP1 and T cells showed that EAP45 is required for efficient HIV budding and spreading (25, 26). The specificity of this effect was demonstrated by the ability of EAP45 *in trans* to rescue the EAP45 KO cell-mediated virus budding and by its dependency on the H0 linker region for interacting with ESCRT-I (26). The discrepancy between these two studies may have arisen from minor amounts of residual ESCRT-II in the siRNA KD study still being sufficient for functional budding. Notably in these studies, cell proliferation was largely unperturbed by the depletion of EAP45 and yet there was up to a 10 fold decrease in virus spreading, suggesting that EAP45 is redundant for cell growth but is critical for viral multiplication and spread (26). This, together with the apparent requirement for early ESCRT proteins to ensure the specific recruitment of the Gag-pol polyprotein (26), underlines the importance of understanding the mechanistic roles of ESCRT-II in HIV budding. Further insights into these protein-protein interactions could help in developing a new class of antivirals to treat HIV. In addition to its link with ESCRT-I in viral budding, the Glue domain in EAP45 contains ubiquitin-binding sites and lipid-binding sites for membrane anchoring at the endosome (27, 28). It was shown in yeast that neither phosphatidylinositol 3-phosphate (PI3P) nor ubiquitin-binding sites are required for endosomal anchoring of EAP45 but the link to ESCRT-I is (22). In metazoans, the Glue domain is believed to be involved in the recruitment of EAP45 at the endosomal membrane, presumably due to its multifaceted roles (28). However, direct visual evidence for this is lacking.

**Fig. 1.**
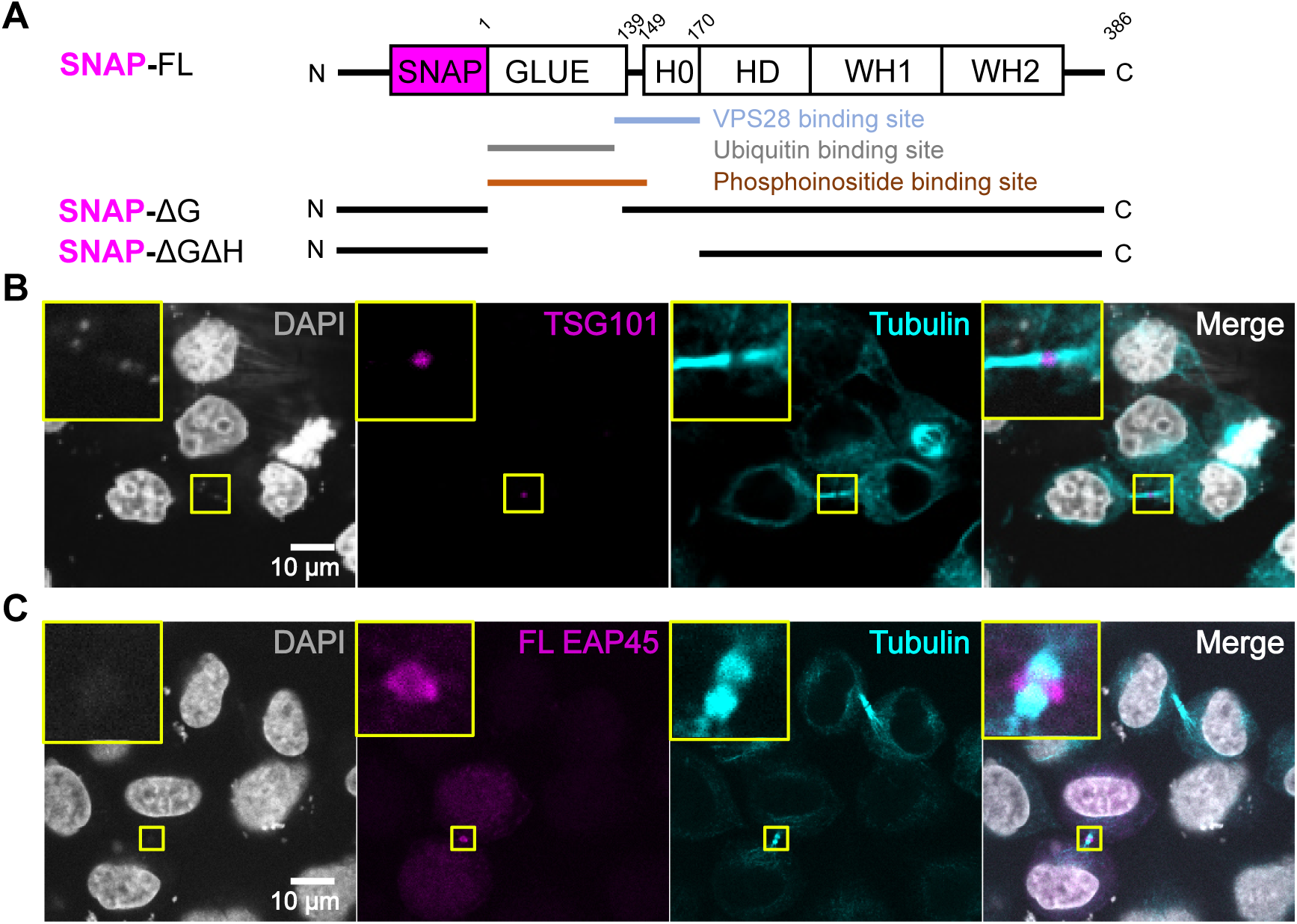
Verification of the SNAP-tag labelling for EAP45 in cytokinesis. **(A)** Schematic diagram showing EAP45 constructs tagged with a SNAP tag at the N terminus and an HA tag at the C terminus (not shown) used in this study. The known interacting domains are highlighted underneath. **(B)** Confocal images of HeLa cells expressing YFP-TSG101 stained with DAPI for nuclei and *α*-tubulin show TSG101 is recruited to the intercellular bridge. **(C)** Confocal images of HeLa cells transfected with SNAP-FL EAP45 followed by staining with DAPI, α-tubulin and Alexa647 conjugated SNAP substrate also show EAP45 is recruited to the intercellular bridge during cytokinesis. The scale is the same for all images.

Building on our previous biochemical evidence supporting the role of ESCRT II in HIV budding, here we used a SNAP-tag to label EAP45 and widefield total internal reflection fluorescence (TIRF) (29) microscopy to visualise and quantify its spatial co-occurrence with Gag polyprotein at the plasma membrane in fixed cells. Furthermore, we used single particle tracking in live HeLa cells to study the dynamics of the interaction between EAP45 and Gag. Lastly, we used this labelling approach to examine the involvement of EAP45 in the endosomal membrane recruitment and during cytokinesis. Our findings show the H0 domain is crucial for anchoring at endosomes and in recruiting ESCRT-I in HIV budding, however the Glue domain is vital in the recruitment of ESCRT-II to the intercellular bridge in cytokinesis.

## Results

### Fusion of a SNAP tag at the N terminus of EAP45 does not affect its normal cellular localisation

The SNAP tag (∼20 kDa) is derived from the mammalian DNA repair protein O^6^-alkylguanine-DNA alkyltransferase and is widely used for imaging studies on both fixed and live cells due to the versatility of the commercially available substrates, providing a superior specificity for targeting the protein of interest (30). A slightly bigger GFP tag (∼27 kDa) was previously inserted at the N terminus of EAP45 without adversely affecting its cellular localisation (28) so we decided to SNAP label EAP45 at the same site (Fig. 1A). Overexpression of ESCRT protein can sometimes have adverse effects on cells (6, 7). To preclude the possibility of the incorporation of a tag affecting the normal cellular localisation of EAP45, we transfected full length (FL) SNAP-EAP45 into HeLa cells and examined the recruitment of EAP45 during cytokinesis, one of the well-characterised activities in which ESCRT proteins are involved (31, 32). A HeLa cell line constitutively expressing YFP-TSG101 was used as a control (31). We observed that TSG101 was recruited to the intercellular bridge as expected (Fig. 1B). Following substrate staining and specificity verification for the SNAP tag (Fig. S1), we also observed that EAP45 was recruited to a similar location (Fig. 1C). The labelling method is specific, as the cells which were not successfully transfected showed no signal from the staining. These data show that the normal cellular activity of EAP45 is not affected by the insertion of an N-terminal SNAP tag.

### SNAP-EAP45 rescues HIV budding in HAP1-EAP45 KO cells

We then verified the rescue functions of the expressor in HIV budding using a similar procedure to that which we reported previously (26). Transient expression of SNAP-FL EAP45 with WT provirus into the HAP1-EAP45 KO cells shows rescue of the virus budding defect (Fig. 2A). This was accompanied by restoration of a normal rate of cleavage at the last step of Gag polyprotein processing (p24/p2), consistent with the effect observed with the non-tagged EAP45 (Fig. 2B) (26). This suggests that fusion of a SNAP tag does not affect the function of EAP45 in rescuing HIV export from HAP1-EAP45 KO cells. To fluorescently label Gag for imaging, we used a piGFP construct in which a GFP tag was inserted between MA and CA of the Gag polyprotein, and has previously been shown to produce GFP-Gag intracellularly (35). To verify this construct in HAP1 EAP45 KO cells, we co-transfected KO cells with piGFP plasmid, VSV-G envelope, and FL EAP45 expressors. Two days post transfection the supernatants were harvested and used to transduce HeLa cells, and then GFP positive cells were counted (Fig. 2C). Our data show a comparable level of GFP expression between EV and EAP45 transfected HAP1 EAP45 KO cells, consistent with the previous observation that expressing EAP45 *in trans* does not alter intracellular viral gene expression (26). However, when the virions were used to transduce HeLa cells, a significantly increased transduction level was observed in EAP45-expressing HAP1 cells in comparison to that of the empty vector (EV). These data confirm that the GFP labelled virus works comparably with the wild-type un-tagged viruses in HAP1 EAP45 KO cells.

**Fig. 2.**
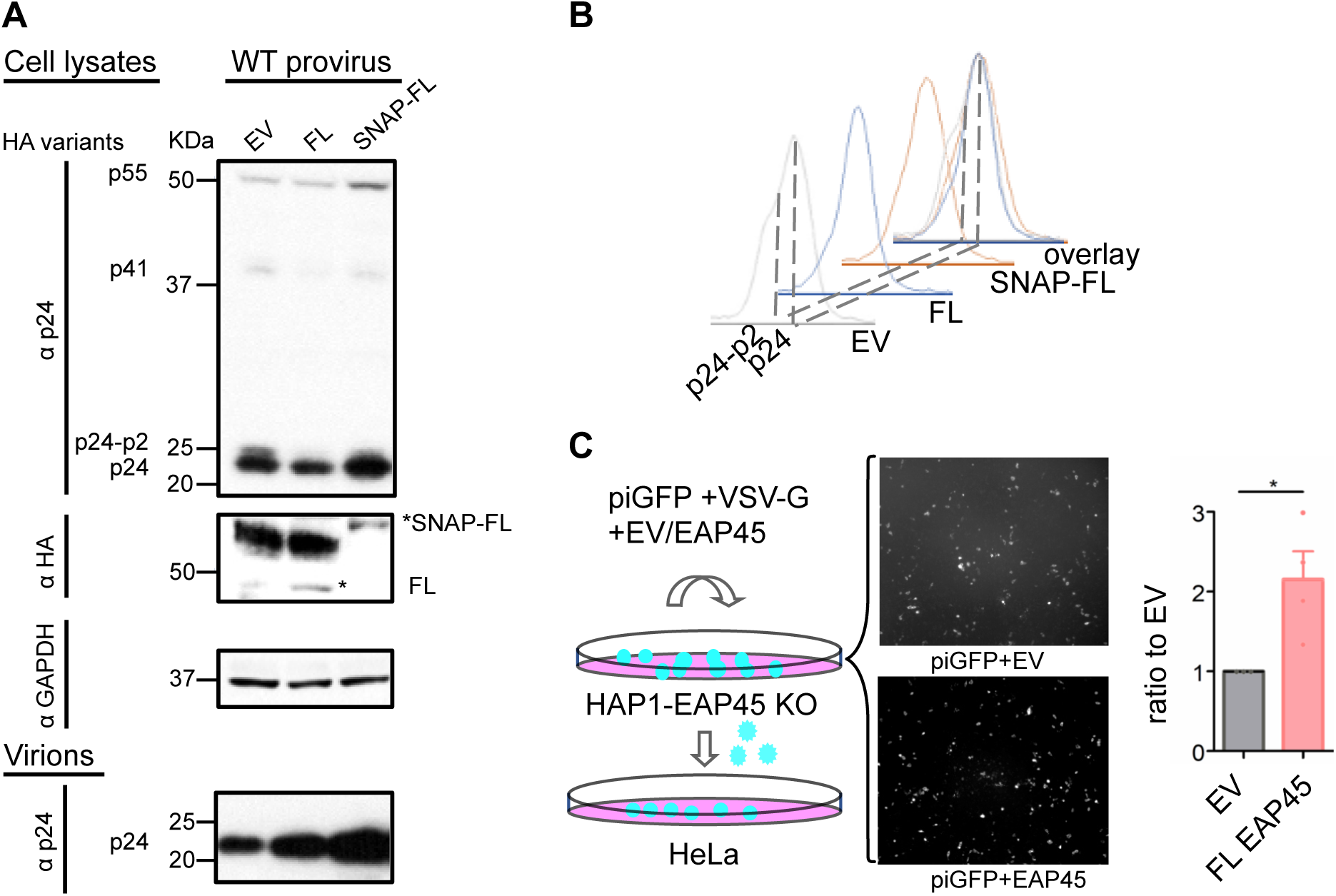
Verification of the SNAP-tag and GFP-Gag in HIV budding in HAP1 EAP45 KO cells. **(A)**The WT provirus was co-transfected with either empty vector (EV), untagged EAP45, or SNAP-FL EAP45 in HAP1 EAP45 KO cells. Cells were lysed two days post transfection and supernatants were harvested for virus purification before western blotting using anti-p24 or anti-HA antibody. GAPDH is also immunoblotted as a control. Protein size markers are indicated by the blot on the left and the densitometric analyses of the final step of Gag cleavage (p24/p24-p2) are shown next to the blots in **(B)**. Asterisks show the band of interest. **(C)** piGFP was co-transfected with EAP45 or EV in HAP1 EAP45 KO cells with fluorescence images taken two days post transfection. The supernatants from the cells were collected and used for transduction onto a layer of HeLa cells for counting GFP positive cells. The number of GFP positive cells was normalised to that of the EV control. The statistical analysis was carried out using a one-sample t test on ≥3 replicates from two experiments. The statistical significance is annotated with p ≤ .05 (*), p ≤ .01 (**), and p ≤ .001 (***) and ns (non-significance) throughout this study.

### Co-occurrence analysis using widefield TIRF microscopy

After verifying the specificity of the SNAP label for EAP45 and the GFP label for Gag, we devised an imaging and analysis procedure to observe and measure the co-occurrence of EAP45 and Gag at the plasma membrane using widefield TIRF microscopy, as illustrated in Fig. 3A. To validate this procedure, we used a nearest-neighbour distance analysis (36, 37) to search for the maximal co-occurrence from two different fluorescent labels on the same species. HeLa cells were transfected with piGFP, followed by staining with an anti-GFP nanobody conjugated with AlexaFluor647 dye. The two different fluorescent markers were expected to perfectly overlap in space, allowing us to test the maximal experimentally attainable colocalisation (Fig. S2A). Images were segmented to detect particles in both colour channels, and the center-to-center distances between Gag and nanobody particles were calculated (Fig. 3B). Any Gag particles found to have a neighbouring nanobody within a given search radius were labelled as “associated”. To account for different densities of Gag particles per cell, the number of associated particles was expressed as a percentage of the total number of detected particles. A search radius of up to 3 pixels (350 nm) was used, and the percentage of associated Gag particles was plotted as a function of this radius (Fig.S2B). The percentage of associated particles at a 1 pixel was ∼75%, increasing to ∼90% within 2 and 3 pixels. These percentages of association provide an estimated upper bound for the maximum proximity expected for a sample in which a marker of interest directly binds to Gag particles. Following validation using this ideal sample, we examined the co-occurrence of Gag and ALIX (Fig. 3C). ALIX is well known to be transiently recruited to the budding site by Gag during viral egress for subsequent recruitment of other ESCRT components (13). We transfected piGFP into an mCherry-ALIX expressing HeLa cell line (31). One day post-transfection, the cells were fixed and examined using TIRF microscopy. The nearest-neighbour analysis procedure described above (Fig. 3B) was applied to search for ALIX particles in the vicinity of Gag particles. Based on the average size of clusters of ES-CRT proteins (∼150 nm) (14), the diameter of HIV budding virions (∼120 nm) (38), and the diffraction-limited resolution of our microscope (2 pixels=234 nm), we selected a 1-pixel perimeter as a constraint for defining interactions between the two clusters. Our data show that around 2.5% of ALIX puncta associated with at least one Gag punctum within a 1-pixel distance (Fig. 3C). This colocalisation rate is similar to the ∼1-4% reported previously for various ESCRT components (15).

**Fig. 3.**
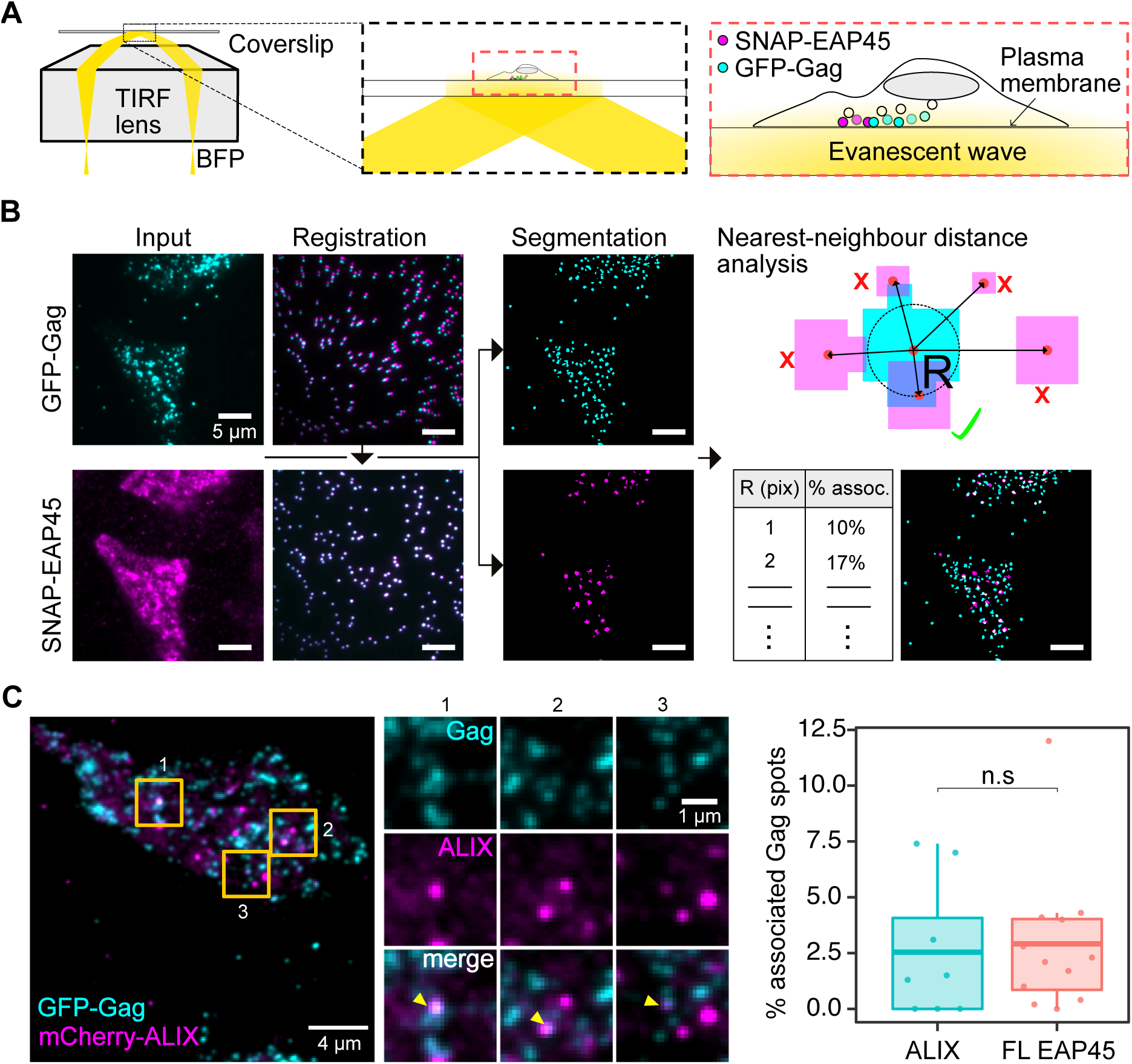
Imaging Gag and EAP45 at the plasma membrane using TIRF microscopy. **(A)** Schematic diagram of the illumination set up used to visualise the spatial proximity of GFP-Gag clusters and EAP45 clusters in mammalian cells. **(B)** Raw dual-colour images captured with the TIRF microscope were registered using calibration images from multicolour fluorescent beads and then segmented using the Trainable Weka Segmentation (33) plugin in ImageJ/Fiji (34). A nearest-neighbour distance analysis was implemented in Matlab to quantify the spatial proximity of Gag and EAP45 or other protein of interest. Particles found within one pixel (∼117 nm) from a particle in the opposite channel were labelled as “associated”. **(C)** Co-occurrence analysis applied to cells expressing ALIX and Gag, as well as cells with SNAP-FL EAP45 and Gag (the same FL EAP45 values in Fig. 4E) for direct comparison). Representative images of the mCherry-ALIX cells with GFP-Gag are shown, along with insets showing instances of co-occurrence between the two markers. Yellow arrowheads in the insets, shown throughout this study, highlight puncta in both channels which show visible co-occurrence. The plot shows the percentage of associated Gag spots. Each data point represents a cell (n = 8 cells for ALIX, and n = 12 cells for FL EAP45) processed with the pipeline illustrated in **(B)**. Welch’s two sample t-test was used to compare the co-occurrence results from ALIX and Gag, to those of FL EAP45 and Gag.

### TIRF microscopy shows high spatial proximity between EAP45 and Gag compared to functionally compromised mutants

Having validated our imaging and co-occurrence analysis procedure, we imaged HeLa cells transfected with piGFP and SNAP-EAP45 followed by staining with an AlexaFluor647-conjugated SNAP substrate (Fig. 4A-E). The cells were fixed at one day post transfection before imaging. We observed both EAP45 and Gag puncta at the plasma membrane, showing that EAP45 is recruited to and/or is active in the proximity of the plasma membrane. This is consistent with the observation made using quantitative EM analysis showing that components of ESCRTs are located at the plasma membrane (39). The nearest-neighbour distance analysis was applied between FL EAP45 and Gag (n=12 cells, Fig. 4A,C,& E) and compared to that of ALIX and Gag (Fig. 3C). We observed around 3% of EAP45 molecules associated with Gag within our 1-pixel threshold similar to the co-occurrence rate observed between ALIX and Gag (Fig. 3C), suggesting that in Hela cells both EAP45 and ALIX are recruited to the budding sites close to the plasma membrane in a similar abundance. The Glue/H0 deletion mutant (ΔGΔH) is deficient at rescuing the HIV budding defect in EAP45 KO cells, as the H0 linker region responsible for this rescue by binding ESCRT-I is absent (26). As expected, removal of the Glue and H0 domains results in lower spatial proximity between EAP45 and Gag particles within a diffraction-limited region than that measured for the FL EAP45 (n = 15 cells, Fig. 4B,D,E, p = 0.026 between FL EAP45 and ΔGΔH), reinforcing the notion that this region plays a pivotal role in linking with Gag. The PTAP domain of Gag p6 is involved in recruiting TSG101 and thus cascading the recruitment of ESCRT-II and ESCRT-III in HIV budding. We hypothesised that the removal of the PTAP late domain (ΔPTAP) would impair the co-occurrence between EAP45 and Gag. Our data however show that even though there is a decrease in the averaged co-occurrence between FL EAP45 with PTAP mutants (n = 11 cells), this is not significantly different from that of WT Gag and FL EAP45 (Fig. 4E, p = 0.36), indicating that EAP45 also co-occurs with Gag independently of PTAP. Consistently, this co-occurrence with the PTAP mutant relies on an intact Glue and H0 linker region, as the removal of this region completely abrogates this phenomenon (n = 14 cells, Fig. 4E, p = 0.018 between FL EAP45 and Δ PTAP-ΔGΔH).

**Fig. 4.**
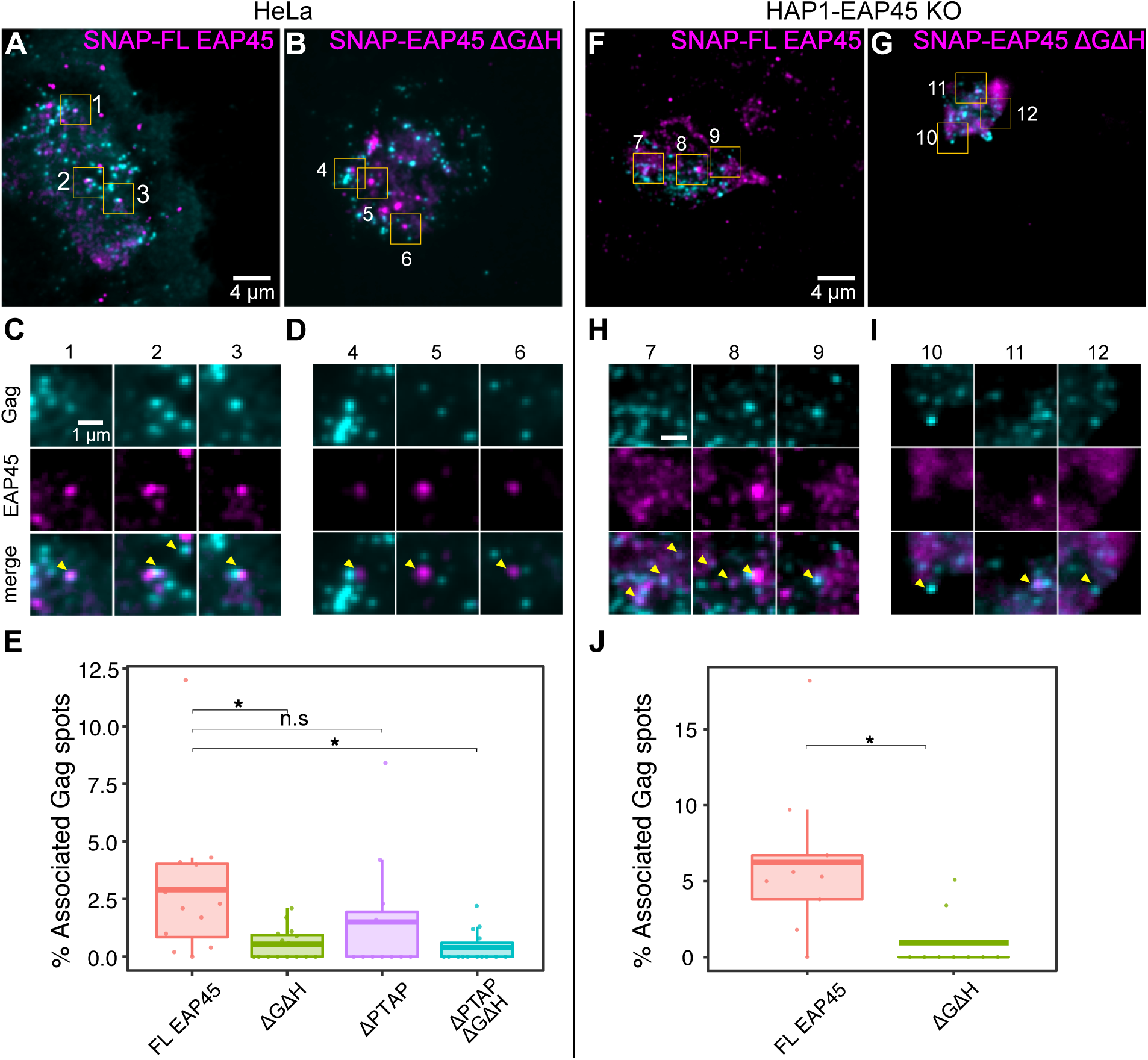
Widefield TIRF microscopy images of fixed HeLa and HAP1-EAP45KO cells with fluorescent markers for Gag and EAP45. **(A)** Representative images of HeLa cells expressing SNAP-FL EAP45 with insets showing regions of high spatial overlap between the Gag and EAP45 markers **(C). (B)** HeLa cells expressing SNAP-ΔGΔH EAP45 with insets showing regions with little spatial overlap between markers **(D)**.**(E)** Comparison of the percentage of associated Gag particles, i.e. those which have an EAP45 particle within 1 pixel, between FL EAP45 (n = 12), ΔGΔH (n = 15), ΔPTAP (n=11), and ΔPTAP-ΔGΔH (n = 14) expressing HeLa cells using a One-Way ANOVA with Dunnett comparisons. **(F)** Representative images for HAP1-EAP45KO cells expressing SNAP-FL EAP45 with insets showing regions of high spatial overlap between the two markers **(H). (G)** Images of HAP1-EAP45KO cells expressing SNAP-ΔGΔH EAP45 with insets showing regions with little spatial overlap between markers **(I). (J)** Comparison of the percentage of associated Gag particles between the FL EAP45 (n = 9) and ΔGΔH EAP45 (n = 9) expressing HAP1-EAP45KO cells using Welch’s t-test.

To further confirm the specificity of the co-occurrence observed between EAP45 and Gag, we also examined them in HAP1-EAP45 KO cells, with subsequent supplementation *in trans* with FL EAP45, followed by image acquisition and analysis (Fig. 4F-J). We tested the proximity of Gag and FL EAP45 (n = 8 cells) versus that of Gag and ΔGΔH EAP45 (n=9 cells). Our data show a significantly higher (p = 0.018) number of EAP45 particles within the association threshold of Gag for FL (mean ∼6%) than for the ΔGΔH mutant (mean ∼1%), as illustrated in Fig. 4J, reinforcing the notion that the N terminus of EAP45 is required for the co-occurrence of EAP45 and Gag by providing a bridge linking EAP45 to ESCRT-I (26). The findings from the nearest-neighbour analysis in both HAP1-EAP45 KO and HeLa cells are consistent with the evidence obtained biochemically showing the N terminus encompassing the H0 linker region in EAP45 plays an important role in HIV budding.

### Single particle tracking in live HeLa cells shows transient co-occurrence of Gag and EAP45

The assembly of Gag particles at the cell membrane during HIV budding has been reported to occur within a time frame of ∼10 mins (10, 40), and the dynamics of ESCRT protein recruitment by Gag particles have been reported for TSG101 (10), VPS4A (5, 11, 14), some CHMP proteins of ESCRT-III (5, 10) and ALIX (13), with an association time at the membrane for different components varying from a few seconds up to 20 mins.

To further validate the observed association between Gag and EAP45, and to analyse its dynamics, we used TIRF microscopy in live HeLa cells to visualise the temporal interaction between GFP-Gag and SNAP-FL EAP45 at the plasma membrane. The HeLa cells were transfected with GFP-Gag and Gag expressors at a 1 : 5 ratio, similarly to previous reports (13, 14), with SNAP-FL EAP45. The substrate for SNAP-EAP45 was added at >10 hours post-transfection when the Gag puncta are predominantly detectable at the plasma membrane (35). We analysed 20-min long live cell movies using single particle tracking. A sub-pixel fitting routine to calculate the centre position of the puncta was applied in which the distance between the centres of an EAP45 particle and a Gag particle were plotted over time. In the live cell movies, both Gag and EAP45 clusters move simultaneously, making it difficult to see how their relative positions are correlated over time. To visualise the relative motion of the particles more clearly, we developed an algorithm that tracked the position of a Gag particle and repositioned the field of view such that the detected particle remained in the centre of the image from frame to frame. We used the Gag particle position as a reference point against which the motion of the EAP45 cluster could be measured. We call this algorithm a “co-moving frame analysis”, and its steps are summarised in Fig. S3. For each region of interest (ROI) extracted, the distance between the Gag and EAP45 tracks was plotted as a function of time. The particles were considered to be “associated” if the distance between them was within the diffraction-limited resolution of our microscope, in this case ∼100 nm (2 pixels). Our analysis shows a transient association, or spatial co-occurrence, between Gag and EAP45 particles over a time scale of 1 to 5 mins, with a mean consecutive association time of ∼1.7 mins (Fig. 5C). This is consistent with the time frame of co-occurrence reported for other ESCRT proteins. Within 14 analysed ROIs (Fig. S4), there appear to be three classes of movements between EAP45 and Gag: i) EAP45 associates with Gag multiple times (Fig. 5A-B top row; 7 out of 14 events); ii) EAP45 approaches Gag and remains associated with Gag for a span of time before leaving again (Fig. 5A-B middle row; 4 out of 14 events); iii) EAP45 approaches Gag and remains with Gag during the time of the recording (Fig. 5A-B bottom row; 3 out of 14 events). Furthermore, for all ROIs analysed, EAP45 particles spent ∼30% of the total observation time within the threshold from a Gag particle (Fig. 5D). It is clear from the distance vs time traces and the overlaid tracks in Fig. 5B that the EAP45 particles transiently slow down, or linger in the vicinity of the Gag particle. This suggests a transient interaction between Gag and EAP45, similar to that of Gag and other ESCRT components during viral egress.

**Fig. 5.**
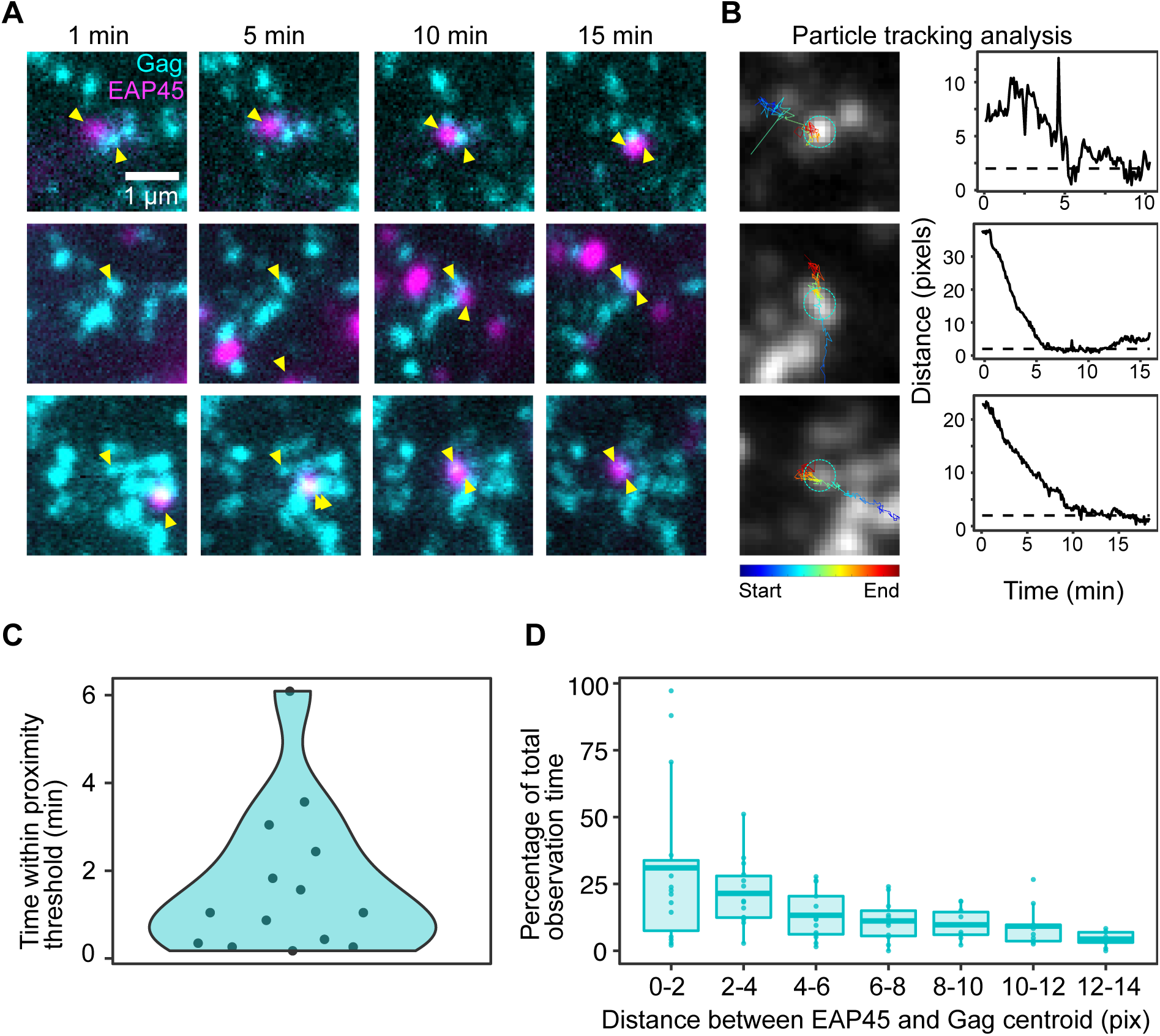
Single particle tracking on live-cell TIRF microscopy images shows a transient co-occurrence between EAP45 and Gag spots from 20 seconds to several minutes. The panels in **(A)** show two-colour images at 1, 5, 10, and 15 mins for different regions of interest in the live acquisitions, with the GFP-Gag channel in cyan and the SNAP-FL EAP45 channel in magenta. Yellow arrowheads are used to point out the virus and EAP45 spots which were selected for single particle tracking. The panels in **(B)** show the results from the co-moving frame analysis. The first column under “Particle tracking analysis” shows an average grey-scale image of the stabilised Gag channel with the selected Gag particle always in the centre of the frame and a super-imposed line plot of the path traced by the EAP45 particle colour-coded for time. The second column under “Particle tracking analysis” shows line plots of the distance in pixels between the centroids of the Gag and EAP45 particles versus time for each of the three regions of interest in **(A)**. The dotted line represents the association threshold of ∼100 nm (2 pixels in the live imaging microscope). **(C)** Kernel density plot showing the maximum consecutive association time between Gag and EAP45 over 14 regions of interest (ROIs). **(D)** Percentage of total observation time as a function of the distance between centroids of Gag and EAP45, plotted over 2-pixel increments (n = 14).

### The Glue domain is not required for the localisation of EAP45 at the late endosome but is required for cytokinesis

ESCRT-II components are also involved in MVB formation and cytokinesis. It has been hypothesised the Glue domain plays an important role in those activities. However, the exact domain requirements of EAP45 remain ill-defined. We used the specific labelling from SNAP tag to investigate this in more detail. EAP45 largely shows a cytosolic distribution and colocalises with RAB7, a late endosome marker, consistent with previous observations (Fig. 6A) (28). Unexpectedly, EAP45 still localises to late endosomes despite removal of the Glue domain (ΔG)(Fig. 6A middle row), suggesting that the Glue domain, which has binding sites for both ubiquitin and lipids, is not required for the recruitment of EAP45 to late endosomes. Further deletion of the H0 linker region (ΔGΔH) greatly reduces the observed colocalisation (Fig. 6A bottom), suggesting the H0 linker region rather than the Glue domain on its own is crucial in anchoring EAP45 at late endosomes. The H0 linker region is required for binding to subunits of ESCRT-I *in vitro* (19, 20), therefore it is possible that ESCRT-I colocalises with and recruits EAP45 at late endosomes. We used a HeLa YFP-TSG101 cell line to examine this possibility. Upon transfection of SNAP-FL EAP45 expressor, we observed a significant cytosolic colocalisation of EAP45 with TSG101 at late endosomes (Fig. 6B). This colocalisation is also observed in the ΔG mutant (Fig. 6B middle) but not in the Δ GΔH mutant (Fig. 6B bottom), suggesting that ESCRT-I plays an important role in the recruitment of ESCRT-II to late endosomes and that this recruitment depends on an intact H0 linker region. In addition to its role in endosomal membrane anchoring, EAP45 is recruited to the intercellular bridge during cytokinesis (Fig. 1C) (41). To better understand which domains in EAP45 are required for this, we transfected expressors of EAP45 and the derived mutants into HeLa cells and imaged the cellular distribution of EAP45 at the intercellular bridge during cytokinesis (Fig. 6C-E). Full length EAP45 is recruited to the intercellular bridge as expected, but this colocalisation is significantly decreased when the Glue domain (ΔG) is removed, suggesting it is important for efficient recruitment in cytokinesis in contrast to the findings in late endosomes. Further removal of the H0 linker region (ΔGΔH) does not significantly decrease recruitment, further stressing the importance of the Glue domain in cytokinesis.

**Fig. 6.**
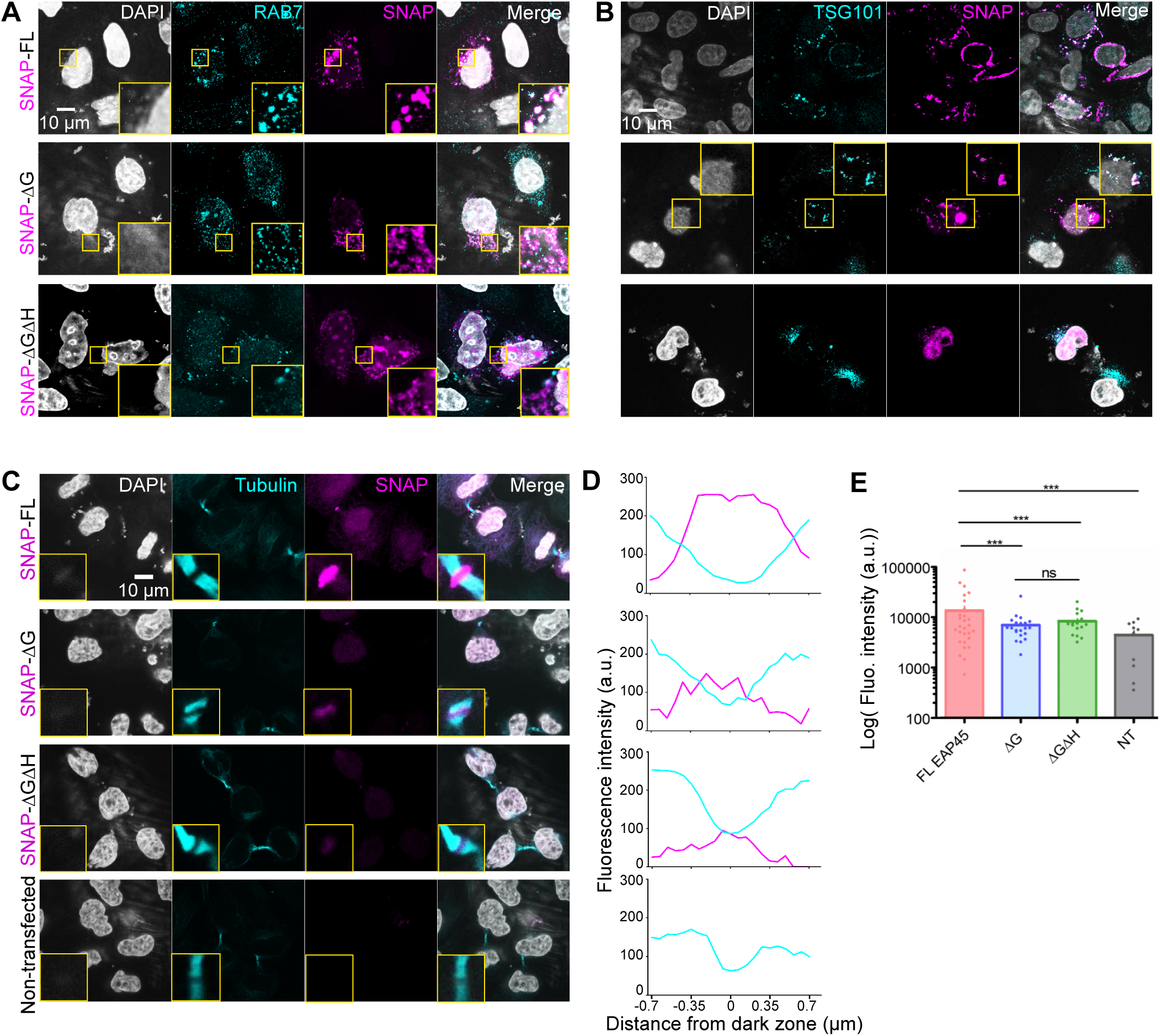
Glue domain is not required for the localisation of EAP45 at the late endosome, but it is required for its localisation at the intercellular bridge during cytokinesis. Confocal images of HeLa cells **(A)** and YFP-TSG101 HeLa cells **(B)** transfected with different SNAP-EAP45 constructs, and stained one day post-transfection with DAPI for nuclei and AlexaFluor647 conjugated SNAP tag substrates for EAP45. The HeLa cells in **(A)** were stained with RAB7, to visualise the colocalisation of late endosomes and EAP45. Close-up views in yellow insets highlight areas of colocalisation. The YFP-TSG101 HeLa cells **(B)** show the colocalisation of TSG101 and EAP45 is similar for the FL and ΔG constructs, but reduced upon deletion of the H0 linker region. **(C)** Confocal images of HeLa cells stained with DAPI, *α*-tubulin, and AlexaFluor647-SNAP substrates for EAP45. **(D)** Plots of line density profiles across the intercellular bridge for each condition shown in **(C)**. **(E)** Quantification of the intensities for each condition where EAP45 is recruited at the intercellular bridge is shown.

## Discussion

The roles of ESCRT-II in MVB formation, cytokinesis, and HIV budding have all been areas of controversy, with previous reports suggesting it is redundant (19, 31, 32). More recent emerging data in cytokinesis (24, 41) and HIV budding (25, 26) appear to revise this and strongly implicate ESCRT-II in a bridging role between ESCRT-I and ESCRT-III as originally reported in cargo sorting (17, 27). Our previous data showed that expressing EAP45 *in trans* rescues an HIV budding defects in HAP1 EAP45 KO cells and biochemical analysis attributed this to the H0 linker domain in EAP45 (26). Using TIRF microscopy we now expand these observations and provide additional clues as to the functions of the different domains of EAP45 in various roles. We observed EAP45 expressed in close proximity to, or at, the plasma membrane in concert with the other ESCRT members documented previously (39). The relatively low co-occurrence rate (around 1 to 6%) between ALIX and Gag (Fig. 3C) and EAP45 and Gag (Fig. 4E & J) is consistent with the rate reported by others using dSTORM to measure both endogenously and exogenously expressed ESCRT components with Gag (1.5 to 3.4%) (15). Considering the maximum experimentally attainable colocalisation measured using the AlexaFluor647 nanobody targeting GFP (Fig. S3), this number is an underestimate of the total events by a factor of 25%. Such a low rate of co-occurrence however is not uncommon (15) and is most likely attributable to the documented transient nature of recruitment of ESCRT proteins. Our data in HAP1 EAP45 KO cells show a higher co-occurrence rate between FL EAP45 and Gag than in HeLa cells (Fig. 4). This highlights the necessity of EAP45’s involvement in HIV budding in HAP1 EAP45 KO cells, since in HeLa cells endogenously expressed but undetected EAP45 could also function *de novo*. Interestingly, the co-occurrence between EAP45 and Gag does not strongly depend on its canonical recruiting motif PTAP, as Δ PTAP Gag reduces the co-occurrence but not to a statistically significant level (Fig. 4E). Given that the level of recruitment and dynamics at the plasma membrane between WT and ΔPTAP Gag are similar (10, 11, 42), this observation could be explained by the recruitment of EAP45 by ALIX via its interaction with TSG101 of ESCRT-I in the absence of the PTAP motif (43). Our co-occurrence analysis between EAP45 and Gag confirms that the N terminal region of EAP45 is crucial in interacting with Gag at the plasma membrane. This is consistent with the biochemical evidence whereby deletion of the Glue and H0 regions completely abolishes the rescue effect mediated by EAP45 in the HAP1 EAP45 KO cells (26), substantiating the importance of this region in linking to ESCRT-I in HIV budding. Interestingly, this interaction is also important in aiding the recruitment of ESCRT-II at the endosomal membrane (Fig. 6B). However, it is less important in the context of ubiquitin and lipid binding, which mirrors previous reports on the recruitment of ESCRT-II at the endosomal membrane in yeast in which H0 mediated anchoring appears more critical than the Glue domain integrity (22). The Glue domain however plays a crucial role in cytokinesis (Fig. 6C-E) possibly by interaction with the unique local lipid components (44). The Glue domain thus unexpectedly plays distinctive roles in different cellular processes.

Our live imaging experiments revealed the dynamic recruitment of EAP45 to the Gag assembling sites (Fig. 5 & 7). On average, EAP45 is within the association distance of Gag particles for a period of 1.7 mins. Remarkably, for 30% of the total recording time (∼5 mins) EAP45 was localised within ∼100 nm of Gag. We observed three distinct classes of movements of EAP45. Some show a similar behaviour to CHMP4B/VPS4 and are recruited multiple times until scission happens (5); others behave similarly to TSG101 in that, once recruited, they serve as an adaptor for the recruitment of downstream components and therefore spend a relatively longer time with Gag (14). The last class in which EAP45 stays with Gag once recruited may be attributable to the molecules that are packaged within the virions, similarly to that observed with ALIX and TSG101 (13, 45). A future study is needed to pinpoint which class of dynamic interaction(s) is necessary and responsible for forming of infectious virus particles as well as the recruitment kinetics of ESCRT-II in the context of other ESCRT proteins.

**Fig. 7.**
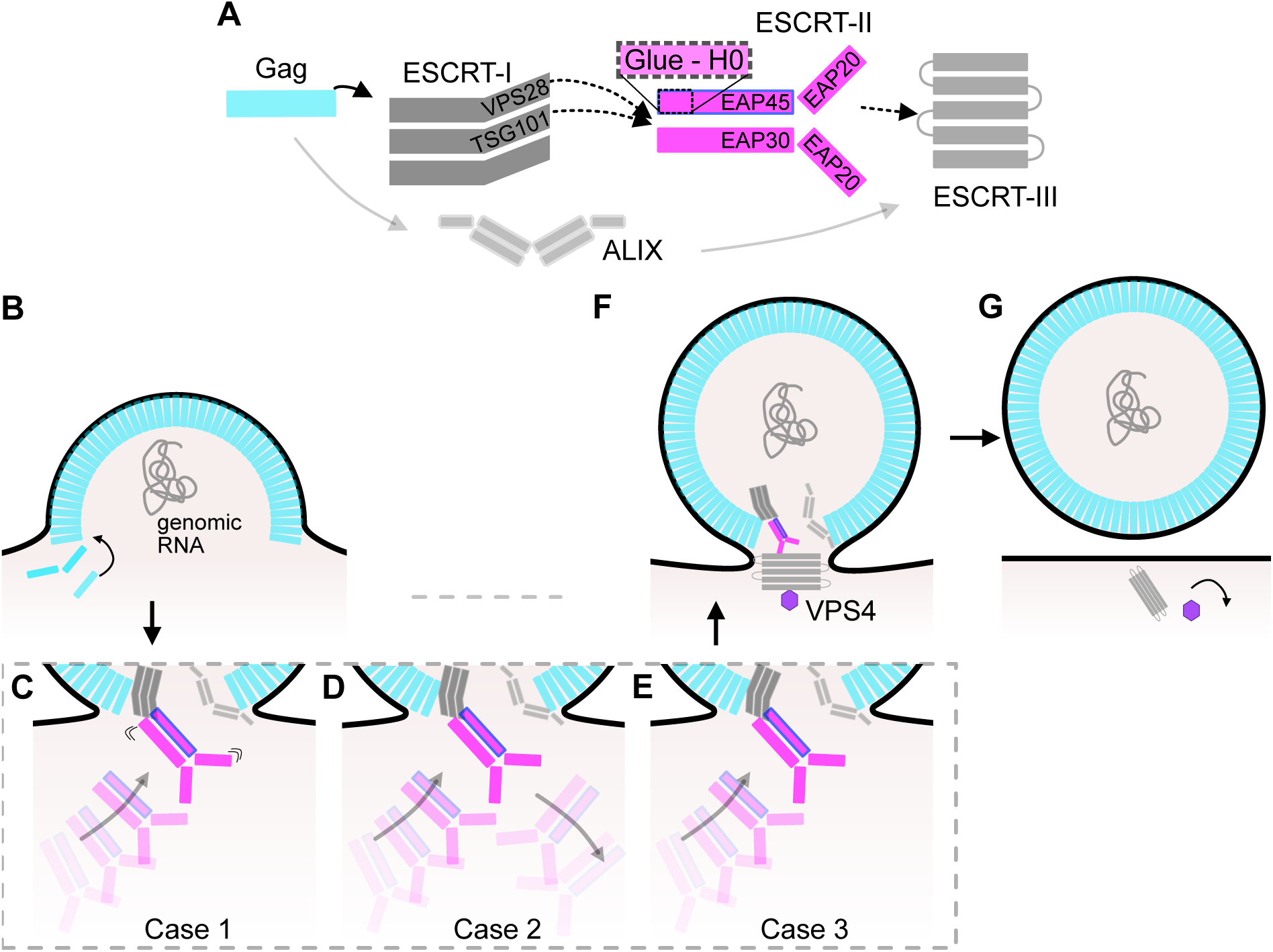
Proposed models for the role of ESCRT-II in HIV budding. **(A)** Cartoon showing the current understanding of the ESCRT protein apparatus in HIV budding. The ESCRT-II complex is shown enlarged, with the EAP45 protein highlighted in blue, and the Glue-H0 region surrounded by dotted lines. **(B)** HIV Gag accumulation causes deformation of the plasma membrane, leading to the recruitment of ESCRTs. Three possible models of ESCRT-II recruitment are illustrated: **(C)** recruitment in which the ESCRT-II oscillates towards and away from the bud, **(D)** transient recruitment followed by decoupling, and **(E)** persistent recruitment. **(F)** shows the final stage before membrane scission in which the polymerised ESCRT-III is recruited to the budding neck and acts in combination with VPS4 to achieve membrane scission and bud egress **(G)**.

In summary, we used a SNAP-tag label to image the main component of ESCRT-II, EAP45, and studied its spatial and temporal co-occurrence with Gag in HIV budding, as well as the domain requirements for its anchoring at the endosomal membrane and in cytokinesis. Our data strengthen the importance of ESCRT-II in linking to ESCRT-I and -III in HIV budding and illustrate similarities to and contrasts with the processes of MVB formation and cytokinesis.

## Materials and Methods

### Plasmids and cells

All cells used in this study were grown in DMEM with 10% FCS at 37 °C in 5% CO2 incubator. HeLa YFP-TSG101 and mCherry-ALIX cells were gifts from Juan Martin-Serrano at KCL. pBH10ΔBglII-WT and pCMV-VSV-G, pEF-EAP45-HA were reported previously (25). piGFP was a gift from Benjamin Chen at Mount Sinai and was described previously (35). A sole Gag expressor (HVPgagpro-) was also described previously (46). SNAP-FL-EAP45, SNAP-ΔG-EAP45 and SNAP-ΔGΔH-EAP45 were constructed by VectorBuilder. The pBH10Gag-GFP-PTAP mutant was cloned by ligating the BssHII and SpeI digested fragments from the piGFP vector to the same sites in pBH10Gag-PTAP, which had been rendered Gag/pol deficient by creating a deletion between BclI and BalI sites.

### Transfection

For transfection, the HeLa cells were transfected with Fugene HD (Promega) and HAP-EAP45 KO cells were transfected with Turbofectin (OriGene). Both the SNAP-expressor rescue experiment and the transfection of piGFP in HAP1-EAP45 KO cells were done similarly as described before (26). Briefly, 150 ng of pBH10ΔBglII-WT or 250 ng of piGFP was co-transfected with 45 ng of SNAP-FL-EAP45 or pEF-EAP45-HA, together with pCMV-VSV-G, respectively, in a well of a 24 well plate. The supernatant was harvested for virion purification and cells were lysed for immunoblotting at two days post transfection. 8-well glass-bottom Lab-Tek slides were used in both fixed and live cell imaging. For fixed cell imaging, HeLa cells were co-transfected with 100 ng piGFP and 60 ng of SNAP-FL-EAP45. HeLa-mCherry-ALIX cells were transfected with 100 ng piGFP. HAP1-EAP45 KO cells were co-transfected with 200 ng piGFP and 72 ng of SNAP-FL-EAP45. At 24 hours post transfection, the cells were fixed, permeabilised and stained with the SNAP-Surface AlexaFluor647 (NEB) at 4 µM for 1 hour. For live cell experiments, a total amount of 120 ng of piGFP and HVPgagproplasmids at a ratio of 1:5 were co-transfected with 72 ng of SNAP-FL-EAP45. At > 10 hours post transfection, the cells were stained with cell permeable live SiR-SNAP (NEB) at 3 µM for 30 mins before examination by widefield TIRF microscopy.

### Immunoblotting

Immunoblotting was done essentially as reported previously (26). Briefly, at 2 days post transfection, supernatants were harvested for virion purification and monolayers were lysed in cell culture lysis buffer (CCLR, Promega) for immunoblotting. The primary antibodies used in this study were anti-p24 (NIBSC), anti-GAPDH (Abcam), anti-HA (Thermofisher), and secondary antibodies used were goat anti-mouse IgG-HRP (Cell Signalling Technology) and goat anti-rabbit IgG-HRP (Thermofisher). Densitometric analysis of the target bands was done in ImageJ (34) with traces of p24 and p24-p2 highlighted to show the differences in the final cleavage product.

### Confocal microscopy

HeLa or HeLa-YFP TSG101 cells were seeded into 8-well slides (EZ slides, Merck). At 24-hour post seeding, the cells were transfected with 100 ng of either SNAP-FL, SNAP-ΔG, or SNAP-ΔGΔH-EAP45 expressor. This was followed by fixation with PFA and staining with SNAP-Surface AlexaFluor647 (NEB) at 4 µM concentration for SNAP-EAP45 and DAPI for nuclei on the following day. A primary antibody to RAB7 (Abcam) or anti-Tubulin (Sigma) was also added on some occasions followed by staining with the secondary antibody conjugated with either AlexaFluor488 or AlexaFluor647 (Thermofisher) before imaging using Leica SP5 confocal fluorescence microscope.

### Widefield TIRF fixed cell imaging

Fixed cell images using widefield TIRF microscopy were acquired with a custom-built microscope based on an Olympus (Center Valley, PA) IX-73 frame with a 488-nm laser (Coherent Sapphire 488-300 CW CDRH), a 561-nm laser (Cobolt Jive 500 561 nm), and a 647-nm laser (MPB Communications Inc. VFL-P-300-647-OEM1-B1). Laser light entering the microscope frame was reflected from a dichroic mirror (Chroma ZT488/561/647rpc) into a 100× 1.49 NA oil objective lens (Olympus UAPON100XOTIRF), and then onto the sample. Light emitted by the sample passed through the dichroic and a set of 25 mm band-pass filters (Semrock FF01-525/45-25, Semrock FF01-600/37-25, and Semrock FF01-680/42-25 respectively for the 488, 561, and 647-nm laser lines) before reaching the microscope side port. Images were then relayed onto two cameras (Andor iXon Ultra 897) by a 1.3x magnification Cairn Twincam image splitter with a dichroic beam splitter (T565spxr-UF3) which allows imaging of emitted wavelengths between 390-555 nm in the transmission port and 575-1000 nm in the reflection port. The pixel size in the sample plane was measured to be 117 nm.

### Nearest-neighbour distance analysis of fixed cell images

We used an object-based nearest-neighbour analysis method in which we segment the registered images for each colour channel, locate the centroid of the fluorescent spots, and search for particles in the opposite colour channel within a neighbourhood of 3 pixels (∼350 nm). A distance threshold within the diffraction-limited resolution of our microscope (1 pixel ∼117 nm) was selected to designate instances in which these molecules could be interacting, and any particles found within this threshold from the centroid of the opposite channel were labelled as “associated”. Raw dual-colour images were registered to correct for chromatic offset using the DoM_Utrecht plugin version 1.1.6 (https://github.com/ekatrukha/DoM_Utrecht) in Fiji (34), using 7 pixels as the maximum shift between images, a value of 15 for the SNR filter, and a PSF standard deviation of 1 pixel. The registered images were then segmented using the Trainable Weka Segmentation plugin from Fiji/ImageJ (33) with the default Fast Random Forest classifier. Three classes were defined (small blobs, large blobs, and background) and images were annotated to identify single fluorescent particles in focus (small blobs), large aggregates of particles (large blobs), and background. The segmented images were then imported into a custom Matlab (The MathWorks, Natick, USA) script which implemented a nearest neighbour algorithm to check, for each identified segmented spot, the distance from its nearest neighbours in the opposite colour channel. The function natsortfiles was used to sort the data (47). All distances for each image were recorded and exported in a .csv file. The .csv files for all experiments were imported into an R (R Core Team, 2013) script that was used to plot the files and run the statistical analysis on the different experimental conditions.

### Widefield TIRF live cell imaging

Live cell images using widefield TIRF microscopy were acquired with a custom-built microscope based on an Olympus (Center Valley, PA) IX-71 frame with a 488-nm laser (Toptica iBeam SMART), a 561-nm laser (Coherent OBIS LS), and a 640-nm laser (Cobolt MLD). Laser light entering the microscope frame was reflected from a dichroic mirror (Chroma ZT405/488/561/640rpc) into a 100x 1.49 NA oil objective lens (Olympus UAPON100XOTIRF), and then onto the sample. Light emitted by the sample passed through the dichroic and a set of 25 mm band-pass filters (Semrock FF01-525/45-25, Semrock FF01-600/37-25, and Semrock FF01-680/42-25 respectively for the 488, 561, and 647-nm laser lines) before reaching the microscope side port. Images were then relayed onto an sCMOS camera (Hamamatsu Orca Flash v4.0). Live cell TIRF images were acquired in two colour channels every 5 or 10 seconds with a 100 ms exposure time over 15-20 minutes. The pixel size at the image plane was is ∼50 nm (6.45 µm pixels at the camera de magnified by the 100x objective and a 1.3x relay lens).

### Single particle tracking in live cell images

The dual-colour live cell movies were registered to correct for chromatic offset using the DoM_Utrecht plugin version 1.1.6 (https://github.com/ekatrukha/DoM_Utrecht) in Fiji (34), and regions of interest of ∼80×80 pixels (∼1.3×1.3 µm^2^) were manually extracted in EAP45 particles were found in the vicinity of a Gag particle. The Gag particles and the EAP45 particles were tracked independently using Track-Mate v4.0.0 (48) using a radius of 4 pixels (∼200 nm), a threshold value of 2 for particle detection with sub-pixel localisation, and the LAP tracker for tracking. The resulting tracks were imported into Matlab for nearest neighbour analysis. A custom Matlab script was written to import Track-Mate tracks. The distances between x,y positions of the tracks in each channel are calculated and output as a .csv file. For ease of visualisation, a new frame is generated centred around the x,y position of the Gag track such that the Gag particle is always at the center of the frame, and the position of the EAP45 particle is shown relative to the new frame of reference. This co-moving frame analysis makes it easy to visualise the motion of two particles over time in a 2D image. We show the average image of the immobilised Gag particle, and the relative motion of EAP45 on top of this image, colour coded as a function of time.

### Statistical Analysis

For the case in which only two experimental data sets were compared, such as in Fig. 3C for ALIX and FL EAP45, and Fig. 4J for FL EAP45 and ΔGΔH, Welch’s two sample t-test was used with ns: non-significance, * for p < 0.05, ** for p <0.01, and *** for p < 0.001. Boxplots were overlaid on the raw data points, with the bold horizontal line showing the mean, and whiskers represent the largest and smallest value within 1.5 times the interquartile range above 75th and 25th percentile, respectively. For the case in which more than two conditions were compared (Fig. 4E), a One-Way ANOVA with Dunnett Comparison was used to compare the FL EAP45 condition to each of the functionally compromised mutants (ΔGΔH, ΔPTAP, ΔPTAP-ΔGΔH) with ns: non-significance, * for p < 0.05, ** for p <0.01, and *** for p < 0.001. All statistical tests were performed using R.

### Data availability statement

The Matlab (Natick, MA) code used for the nearest-neighbour analysis for the fixed cell co-occurrence experiments, and the co-moving frame analysis for the live cell tracking experiments are available at our online repository. The R scripts used to generate the plots in Fig. 3, 4, 5, S1, S2, and S4 can also be found in this repository.

## ACKNOWLEDGEMENTS

We would like to thank Eric J. Rees and John Sinclair for helpful discussions. We are grateful to Juan Martin-Serrano and Benjamin Chen for providing the cell lines and plasmid, respectively, used in this study. We also thank Ricardo Henriques for his BioRxiv preprint LateX template. This work is supported by the Research Visit Grant by the Microbiology Society (BM), Gates Cambridge Scholarship (PVR), UK Medical Research Council (MRC) Grant (MR/N0229939/1)(JCK and AMLL), MRC Grants (MR/ K015850/1 and MR/K02292X/1)(CFK), UK Engineering and Physical Sciences Research Council, EPSRC Grants (EP/L015889/1 and EP/H018301/1)(CFK), Wellcome Trust Grants (203249/Z/16/Z and 089703/Z/09/Z)(CFK), MedImmune, the RCUK under the Technology Touching Life Initiative, and Infinitus China Ltd (CFK). We also acknowledge the support from the Cambridge NIHR BRC Cell Phenotyping Hub on confocal imaging.

## AUTHOR CONTRIBUTIONS

**Bo Meng**,Conceptualization, Funding Acquisition, Data Curation, Formal analysis, Investigation, Methodology, Validation, Visualization, Writing-original draft, Writing-review and editing; **Pedro P. Vallejo Ramirez**, Data Curation, Formal analysis, Investigation, Methodology, Software, Validation, Visualization, Writing-original draft, Writing-review and editing; **Katharina Scherer**, Data Curation, Investigation, Writing-review and editing; **Ezra Bruggeman**, Software; **Julia C. Kenyon**, Conceptualization, Funding acquisition, Project Administration, Supervision, Writing-review and editing; **Clemens F. Kaminski**, Conceptualization, Funding acquisition, Project Administration, Resources, Supervision, Writing-review and editing; **Andrew M. Lever**, Conceptualization, Funding acquisition, Project Administration, Resources, Supervision, Writing-review and editing.

---

**Fig. S1.**
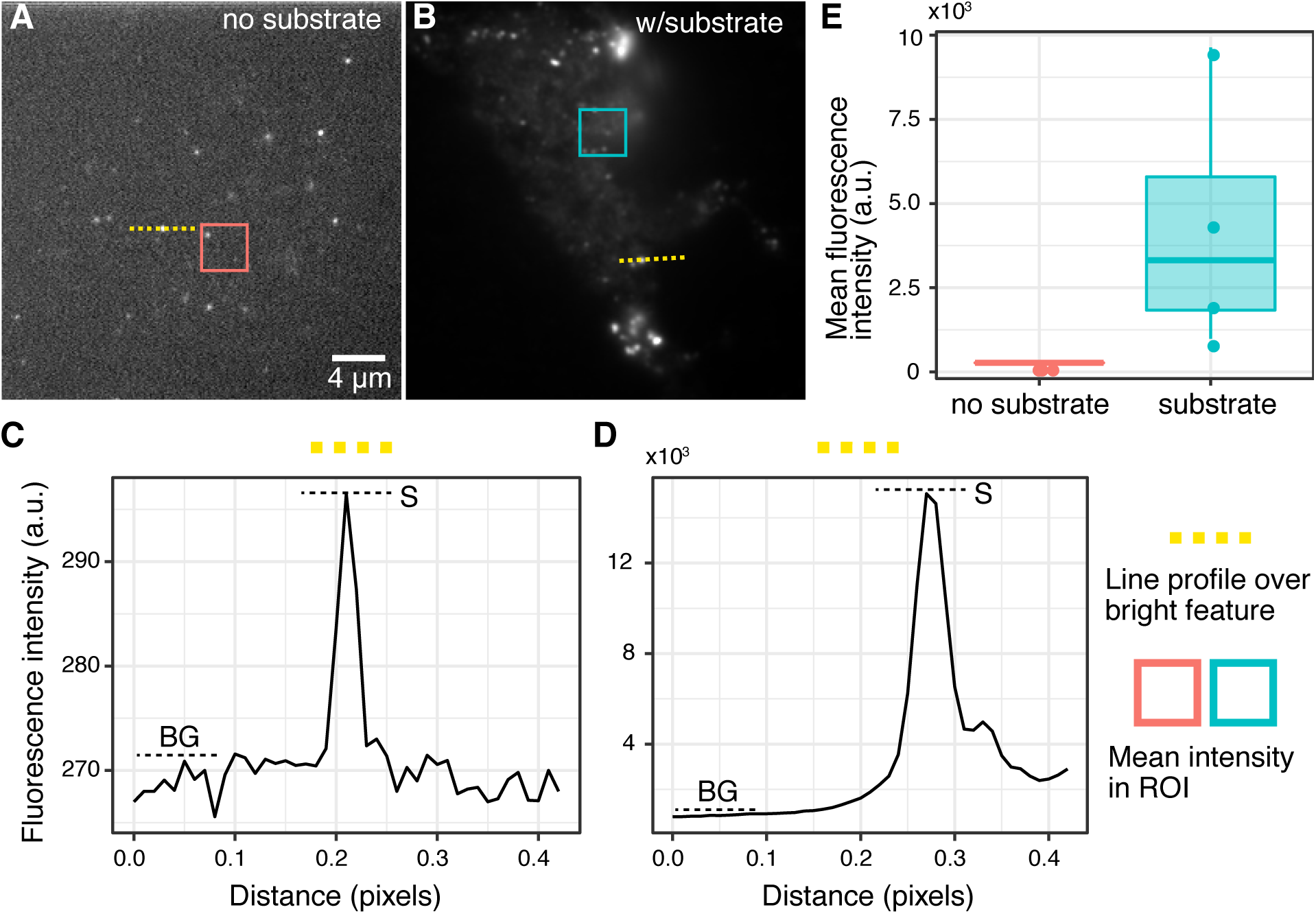
Verification of the labelling strategy using SNAP-tag for EAP45. **(A)** shows an image of HeLa cells with no addition of the SNAP substrate, along with a line profile taken across a bright feature and plotted in **(C)** showing the signal intensity is predominantly background with a mean level around 270 a.u., and the signal to background ratio (SBR) is 1.1. **(B)** shows an image of a cell with the addition of the SNAP substrate in which the mean signal level on a line profile through a bright feature plotted in **(D)** is 2000 a.u., and the SBR is 7, a sevenfold increase from **(B). (E)** shows a comparison of the mean signal inside a square region of interest taken in the absence or presence of the substrate, to indicate the specificity on the labelling (n=4 cells). The contrast levels in image **(A)** and **(B)** are different to visualise the much lower intensity features in **(A)**. The scale is the same for both images.

**Fig. S2.**
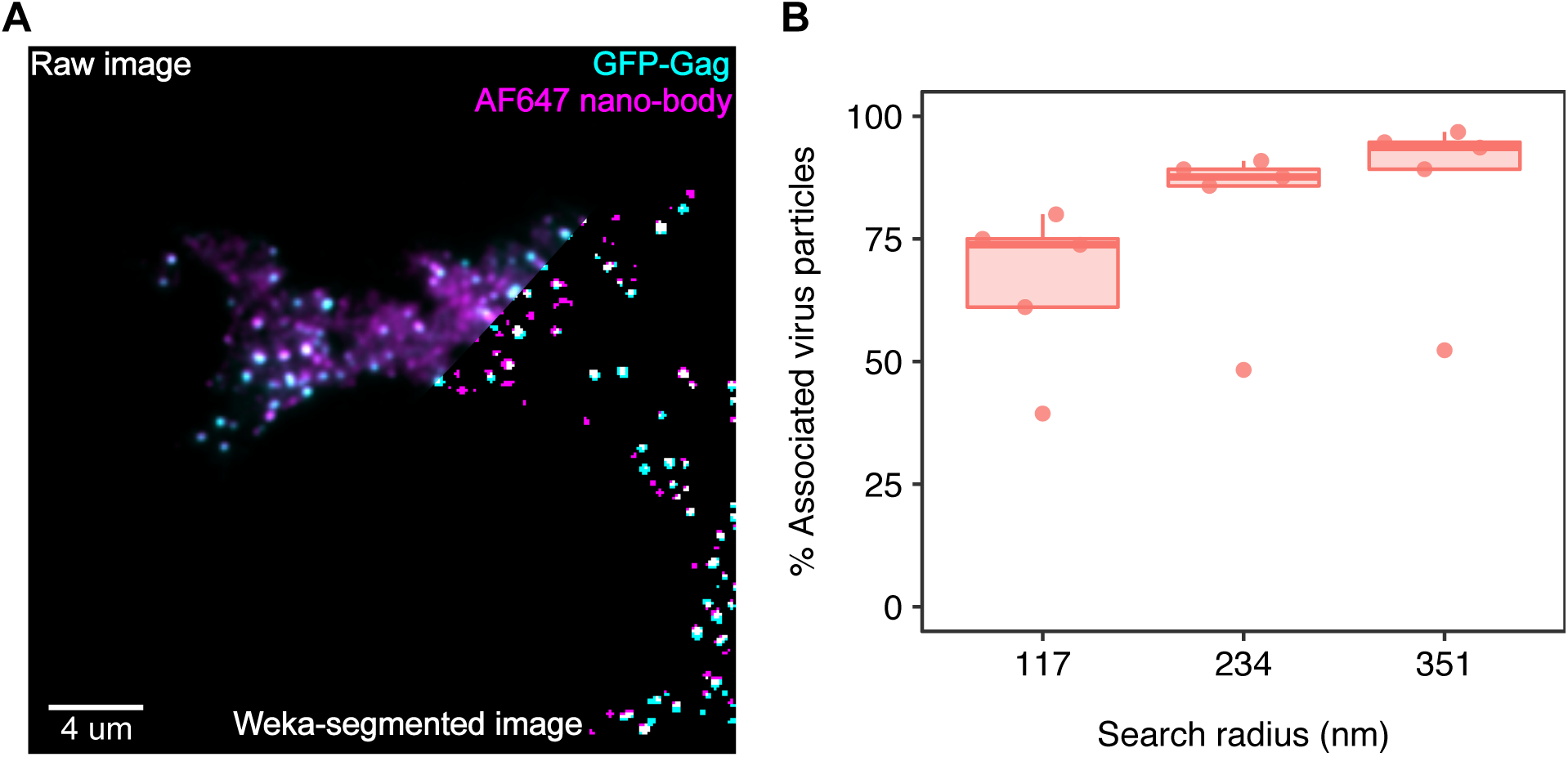
TIRF image of GFP-Gag and anti-GFP nanobody conjugated AlexaFluor647 to test the maximal experimentally attainable colocalisation. **(A)** shows a representative dual-colour image of GFP-Gag and the AlexaFluor647 nanobody, with a faded-in composite towards the lower right corner showing the resulting dual-colour mask after Weka segmentation (1). Panel **(B)**(n=5 cells) displays the percentage of Gag particles with at least one AlexaFluor647 particle in proximity within a given radius, as a function of the search radius.

**Fig. S3.**
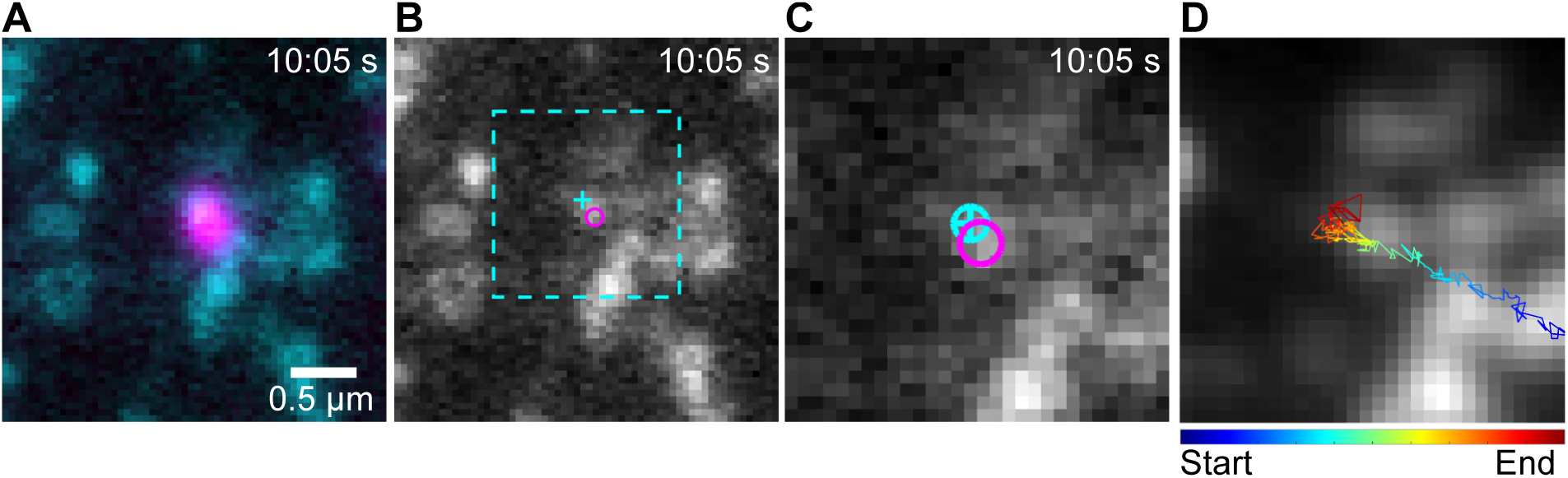
Co-moving frame analysis for single particle tracking data. **(A)** Registered dual colour still frame from live cell movie showing a single EAP45 particle in the vicinity of a Gag particle. **(B)** The centre position of the tracks for a Gag particle (cyan cross) and EAP45 particle (magenta circle) overlaid on a grey scale still frame of the Gag channel. A dotted rectangle indicates a region around the centre position of the Gag channel which is extracted in each frame to immobilise the Gag particle, and the position of the EAP45 molecule is then plotted relative to the new frame of reference in **(C). (D)** shows the track of the EAP45 particle motion colour coded from start to finish of the track duration, overlaid on an average image of the Gag channel.

**Fig. S4.**
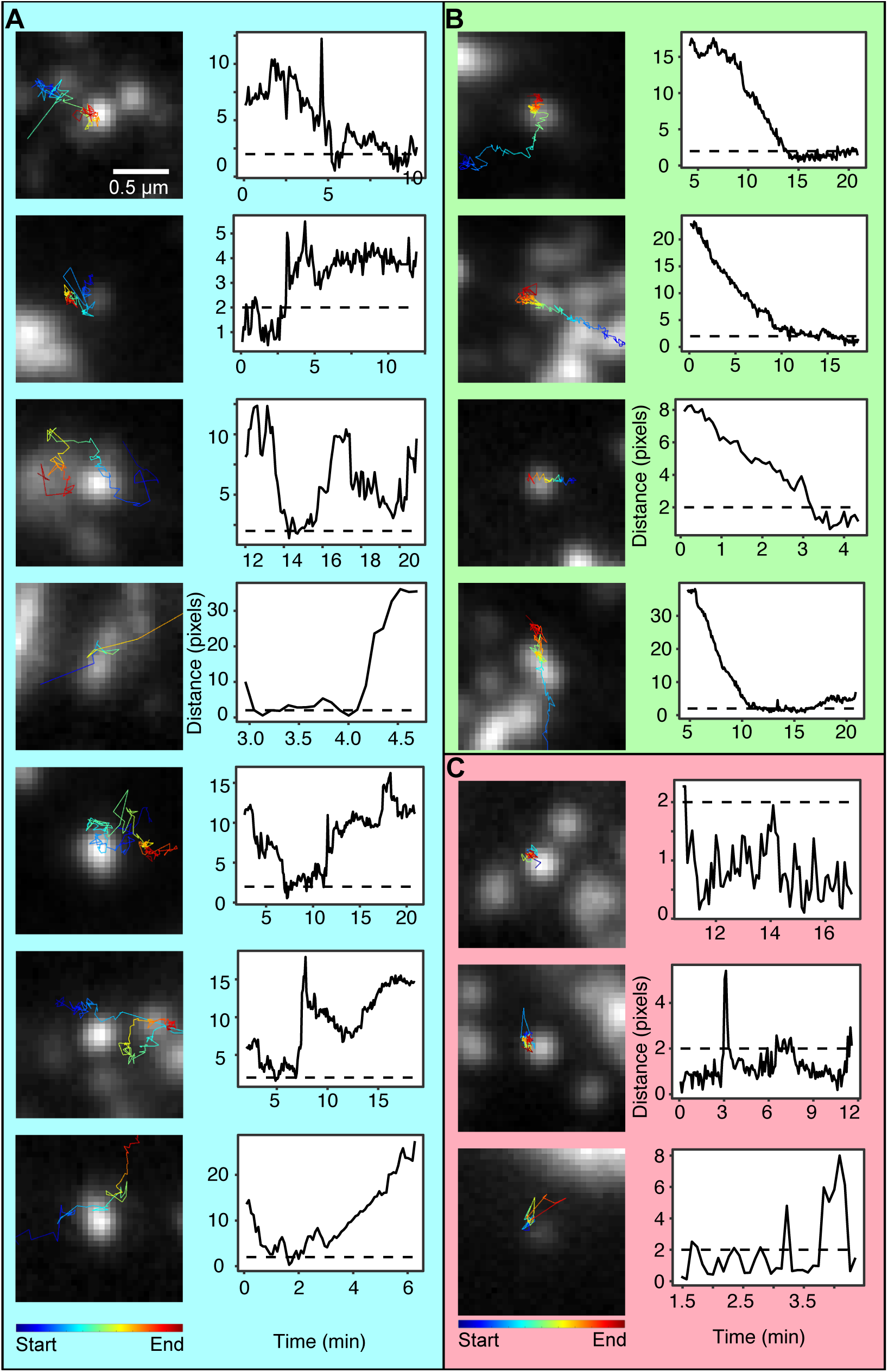
Co-moving frame analysis reveals three classes of EAP45 movement relative to Gag. The motion of the EAP45 relative to the Gag cluster can be classified into three categories: (**(A)**, cyan box) EAP45 particles which hover in the vicinity of the Gag cluster for short spans of time, and either oscillate in and out of proximity, or move away from the Gag cluster, (**(B)**, green box) EAP45 particles which approach the virus from a distance and remain in the 2-pixel vicinity of the virus particle for several minutes, and (**(C)**, red box) EAP45 particles which remain within 2 pixels of the virus particle throughout the observation time interval.

## Bibliography

1. Roger L. Williams and Sylvie Urbé. The emerging shape of the ESCRT machinery. Nature Reviews Molecular Cell Biology, 8(5):355–368, 2007. ISSN 14710072. doi: 10.1038/nrm2162.

2. Johannes Schöneberg, Mark Remec Pavlin, Shannon Yan, Maurizio Righini, Il Hyung Lee, Lars Anders Carlson, Amir Houshang Bahrami, Daniel H. Goldman, Xuefeng Ren, Gerhard Hummer, Carlos Bustamante, and James H. Hurley. ATP-dependent force generation and membrane scission by ESCRT-III and Vps4. Science, 362(6421):1423–1428, 2018. ISSN 10959203. doi: 10.1126/science.aat1839.

3. Wesley I Sundquist and Hans-Georg Krausslich. HIV-1 Assembly, Budding, and Maturation. Cold Spring Harbor Perspectives in Medicine, pages 1–24, 2012.

4. Bo Meng and Andrew M.L. Lever. Wrapping up the bad news - HIV assembly and release. Retrovirology, 10(1):1, 2013. ISSN 17424690. doi: 10.1186/1742-4690-10-5.

5. Daniel S. Johnson, Marina Bleck, and Sanford M. Simon. Timing of ESCRT-III protein recruitment and membrane scission during HIV-1 assembly. eLife, 7:1–20, 2018. ISSN 2050084X. doi: 10.7554/eLife.36221.

6. Bettina Strack, Arianna Calistri, Stewart Craig, Elena Popova, and Heinrich G. Göttlinger. AIP1/ALIX is a binding partner for HIV-1 p6 and EIAV p9 functioning in virus budding. Cell, 114(6):689–699, 2003. ISSN 00928674. doi: 10.1016/S0092-8674(03)00653-6.

7. Uta K. Von Schwedler, Melissa Stuchell, Barbara Müller, Diane M. Ward, Hyo Young Chung, Eiji Morita, Hubert E. Wang, Thaylon Davis, Gong Ping He, Daniel M. Cimbora, Anna Scott, Hans Georg Kräusslich, Jerry Kaplan, Scott G. Morham, and Wesley I. Sundquist. The protein network of HIV budding. Cell, 114(6):701–713, 2003. ISSN 00928674. doi: 10.1016/S0092-8674(03)00714-1.

8. Juan Martin-Serrano and Paul. D. Bieniasz. A Bipartite Late-Budding Domain in Human Immunodeficiency Virus Type 1. Journal of Virology, 77(22):12373–12377, 2003. ISSN 0022-538X. doi: 10.1128/jvi.77.22.12373-12377.2003.

9. Dimiter G. Demirov, Jan M. Orenstein, and Eric O. Freed. The Late Domain of Human Immunodeficiency Virus Type 1 p6 Promotes Virus Release in a Cell Type-Dependent Manner. Journal of Virology, 76(1):105–117, 2002. ISSN 0022-538X. doi: 10.1128/jvi.76.1.105-117.2002.

10. Nolwenn Jouvenet, Maria Zhadina, Paul D Bieniasz, and Sanford M Simon. Dynamics of ESCRT protein recruitment during retroviral assembly. Nature Cell Biology, 13(4):394–401, 2011. ISSN 1465-7392. doi: 10.1038/ncb2207.

11. Viola Baumgärtel, Sergey Ivanchenko, Aurélie Dupont, Mikhail Sergeev, Paul W Wiseman, Hans Georg Kräusslich, Christoph Bräuchle, Barbara Müller, and Don C Lamb. Live-cell visualization of dynamics of HIV budding site interactions with an ESCRT component. Nature Cell Biology, 13(4):469–476, 2011. ISSN 14657392. doi: 10.1038/ncb2215.

12. Sergey Ivanchenko, William J Godinez, Marko Lampe, Hans-Georg Kräusslich, Roland Eils, Karl Rohr, Christoph Bräuchle, Barbara Müller, and Don C Lamb. Dynamics of HIV-1 Assembly and Release. PLOS Pathogens, 5(11):e1000652, nov 2009.

13. Pei I. Ku, Mourad Bendjennat, Jeff Ballew, Michael B. Landesman, and Saveez Saffarian. ALIX is recruited temporarily into HIV-1 budding sites at the end of Gag assembly. PLoS ONE, 9(5):1–8, 2014. ISSN 19326203. doi: 10.1371/journal.pone.0096950.

14. Marina Bleck, Michelle S. Itano, Daniel. S. Johnson, V. Kaye Thomas, Alison J. North, Paul D. Bieniasz, and Sanford M. Simon. Temporal and spatial organization of ESCRT protein recruitment during HIV-1 budding. Proceedings of the National Academy of Sciences, 111(33):12211–12216, 2014. ISSN 0027-8424. doi: 10.1073/pnas.1321655111.

15. Jens Prescher, Viola Baumgärtel, Sergey Ivanchenko, Adriano A Torrano, Christoph Bräuchle, Barbara Müller, and Don C. Lamb. Super-Resolution Imaging of ESCRT-Proteins at HIV-1 Assembly Sites. PLoS Pathogens, 11(2):1–26, 2015. ISSN 15537374. doi: 10.1371/journal.ppat.1004677.

16. Edward J. Scourfield and Juan Martin-Serrano. Growing functions of the ESCRT machinery in cell biology and viral replication. Biochemical Society Transactions, 45(3):613–634, 2017. ISSN 0300-5127. doi: 10.1042/bst20160479.

17. Hsiangling Teo, Olga Perisic, Beatriz González, and Roger L. Williams. ESCRT-II, an endosome-associated complex required for protein sorting: Crystal structure and interactions with ESCRT-III and membranes. Developmental Cell, 7(4):559–569, 2004. ISSN 15345807. doi: 10.1016/j.devcel.2004.09.003.

18. David J. Gill, Hsiangling Teo, Ji Sun, Olga Perisic, Dmitry B. Veprintsev, Scott D. Emr, and Roger L. Williams. Structural insight into the ESCRT-I/-II link and its role in MVB trafficking. EMBO Journal, 26(2):600–612, 2007. ISSN 02614189. doi: 10.1038/sj.emboj.7601501.

19. Charles Langelier, Uta K. von Schwedler, Robert D. Fisher, Ivana De Domenico, Paul L. White, Christopher P. Hill, Jerry Kaplan, Diane Ward, and Wesley I. Sundquist. Human ESCRT-II Complex and Its Role in Human Immunodeficiency Virus Type 1 Release. Journal of Virology, 80(19):9465–9480, 2006. ISSN 0022-538X. doi: 10.1128/jvi.01049-06.

20. Young Jun Im and James H. Hurley. Integrated Structural Model and Membrane Targeting Mechanism of the Human ESCRT-II Complex. Developmental Cell, 14(6):902–913, 2008. ISSN 15345807. doi: 10.1016/j.devcel.2008.04.004.

21. Estela Pineda-Molina, Hassan Belrhali, Andrew J. Piefer, Indira Akula, Paul Bates, and Winfried Weissenhorn. The crystal structure of the C-terminal domain of Vps28 reveals a conserved surface required for Vps20 recruitment. Traffic, 7(8):1007–1016, 2006. ISSN 13989219. doi: 10.1111/j.1600-0854.2006.00440.x.

22. Shrawan Kumar Mageswaran, Natalie K. Johnson, Greg Odorizzi, and Markus Babst. Constitutively active ESCRT-II suppresses the MVB-sorting phenotype of ESCRT-0 and ESCRT-I mutants. Molecular Biology of the Cell, 26(3):554–568, 2015. ISSN 19394586. doi: 10.1091/mbc.E14-10-1469.

23. Shaogeng Tang, Nicholas J. Buchkovich, W. Mike Henne, Sudeep Banjade, Yun Jung Kim, and Scott D. Emr. ESCRT-III activation by parallel action of ESCRT-I/II and ESCRT-0/Bro1 during MVB biogenesis. eLife, 5(April 2016):1–12, 2016. ISSN 2050084X. doi: 10.7554/eLife.15507.

24. Liliane Christ, Eva M. Wenzel, Knut Liestøl, Camilla Raiborg, Coen Campsteijn, and Harald Stenmark. ALIX and ESC RT-I/II function as parallel ESC RT-III recruiters in cytokinetic abscission. Journal of Cell Biology, 212(5):499–513, 2016. ISSN 15408140. doi: 10.1083/jcb.201507009.

25. Bo Meng, Natasha C.Y. Ip, Liam J. Prestwood, Truus E.M. Abbink, and Andrew M.L. Lever. Evidence that the endosomal sorting complex required for transport-II (ESCRT-II) is required for efficient human immunodeficiency virus-1 (HIV-1) production. Retrovirology, 12(1):1–15, 2015. ISSN 17424690. doi: 10.1186/s12977-015-0197-x.

26. Bo Meng, Natasha C.Y. Ip, Truus E.M. Abbink, Julia C. Kenyon, and Andrew M.L. Lever. ESCRT-II functions by linking to ESCRT-I in human immunodeficiency virus-1 budding. Cellular Microbiology, (June 2019):1–15, 2020. ISSN 14625822. doi: 10.1111/cmi.13161.

27. Hsiangling Teo, David J. Gill, Ji Sun, Olga Perisic, Dmitry B. Veprintsev, Yvonne Vallis, Scott D. Emr, and Roger L. Williams. ESCRT-I Core and ESCRT-II GLUE Domain Structures Reveal Role for GLUE in Linking to ESCRT-I and Membranes. Cell, 125(1):99–111, 2006. ISSN 00928674. doi: 10.1016/j.cell.2006.01.047.

28. Thomas Slagsvold, Rein Aasland, Satoshi Hirano, Kristi G. Bache, Camilla Raiborg, Daniel Trambaiolo, Soichi Wakatsuki, and Harald Stenmark. Eap45 in mammalian ESCRT-II binds ubiquitin via a phosphoinositide-interacting GLUE domain. Journal of Biological Chemistry, 280(20):19600–19606, 2005. ISSN 00219258. doi: 10.1074/jbc.M501510200.

29. Daniel Axelrod. Total Internal Reflection Fluorescence Microscopy. Traffic, 2:764–774, 2001. doi: 10.1016/B978-0-12-394447-4.20089-8.

30. Antje Keppler, Maik Kindermann, Susanne Gendreizig, Horst Pick, Horst Vogel, and Kai Johnsson. Labeling of fusion proteins of O6-alkylguanine-DNA alkyltransferase with small molecules in vivo and in vitro. Methods, 32(4):437–444, 2004. ISSN 10462023. doi: 10.1016/j.ymeth.2003.10.007.

31. Jez G. Carlton and Juan Martin-Serrano. Parallels between cytokinesis and retroviral budding: A role for the ESCRT machinery. Science, 316(5833):1908–1912, 2007. ISSN 00368075. doi: 10.1126/science.1143422.

32. Eiji Morita, Virginie Sandrin, Hyo Young Chung, Scott G. Morham, Steven P. Gygi, Christopher K. Rodesch, and Wesley I. Sundquist. Human ESCRT and ALIX proteins interact with proteins of the midbody and function in cytokinesis. EMBO Journal, 26(19):4215–4227, 2007. ISSN 14602075. doi: 10.1038/sj.emboj.7601850.

33. Ignacio Arganda-Carreras, Verena Kaynig, Curtis Rueden, Kevin W Eliceiri, Johannes Schindelin, Albert Cardona, and H Sebastian Seung. Trainable Weka Segmentation : a machine learning tool for microscopy pixel classification. Bioinformatics, 33(March):2424–2426, 2017. doi: 10.1093/bioinformatics/btx180.

34. Johannes Schindelin, Ignacio Arganda-Carreras, Erwin Frise, Verena Kaynig, Mark Longair, Tobias Pietzsch, Stephan Preibisch, Curtis Rueden, Stephan Saalfeld, Benjamin Schmid, Jean-Yves Tinevez, Daniel James White, Volker Hartenstein, Kevin Eliceiri, Pavel Tomancak, and Albert Cardona. Fiji: an open-source platform for biological-image analysis. Nat Meth, 9(7):676–682, jul 2012. ISSN 1548-7091.

35. Wolfgang Hubner, Ping Chen, Armando Del Portillo, Yuxin Liu, Ronald E. Gordon, and Benjamin K. Chen. Sequence of Human Immunodeficiency Virus Type 1 (HIV-1) Gag Localization and Oligomerization Monitored with Live Confocal Imaging of a Replication-Competent, Fluorescently Tagged HIV-1. Journal of Virology, 81(22):12596–12607, 2007. ISSN 0022-538X. doi: 10.1128/jvi.01088-07.

36. E. Lachmanovich, D. E. Shvartsman, Y. Malka, C. Botvin, Y. I. Henis, and A. M. Weiss. Co-localization analysis of complex formation among membrane proteins by computerized fluorescence microscopy: Application to immunofluorescence co-patching studies. Journal of Microscopy, 212(2):122–131, 2003. ISSN 00222720. doi: 10.1046/j.1365-2818.2003.01239.x.

37. Thibault Lagache, Nathalie Sauvonnet, Lydia Danglot, and Jean Christophe Olivo-Marin. Statistical analysis of molecule colocalization in bioimaging. Cytometry Part A, 87(6):568–579, 2015. ISSN 15524930. doi: 10.1002/cyto.a.22629.

38. John A.G. Briggs, Kay Grünewald, Bärbel Glass, Friedrich Förster, Hans Georg Kräusslich, and Stephen D. Fuller. The mechanism of HIV-1 core assembly: Insights from three-dimensional reconstructions of authentic virions. Structure, 14(1):15–20, 2006. ISSN 09692126. doi: 10.1016/j.str.2005.09.010.

39. Sonja Welsch, Anja Habermann, Stefanie Jäger, Barbara Müller, Jacomine Krijnse-Locker, and Hans Georg Kräusslich. Ultrastructural analysis of ESCRT proteins suggests a role for endosome-associated tubular-vesicular membranes in ESCRT function. Traffic, 7(11): 1551–1566, 2006. ISSN 13989219. doi: 10.1111/j.1600-0854.2006.00489.x.

40. Nolwenn Jouvenet, Paul D Bieniasz, and Sanford M Simon. Imaging the biogenesis of individual HIV-1 virions in live cells. Nature, 454(7201):236–240, 2008. ISSN 14764687. doi: 10.1038/nature06998.

41. Inna Goliand, Dikla Nachmias, Ofir Gershony, and Natalie Elia. Inhibition of ESCRT-II-CHMP6 interactions impedes cytokinetic abscission and leads to cell death. Molecular Biology of the Cell, 25(23):3740–3748, 2014. ISSN 19394586. doi: 10.1091/mbc.E14-08-1317.

42. Sebla B. Kutluay, Trinity Zang, Daniel Blanco-Melo, Chelsea Powell, David Jannain, Manel Errando, and Paul D. Bieniasz. Global changes in the RNA binding specificity of HIV-1 gag regulate virion genesis. Cell, 159(5):1096–1109, 2014. ISSN 10974172. doi: 10.1016/j.cell.2014.09.057.

43. Xi Zhou, Jiali Si, Joe Corvera, Gary E. Gallick, and Jian Kuang. Decoding the intrinsic mechanism that prohibits ALIX interaction with ESCRT and viral proteins. Biochemical Journal, 432(3):525–534, 2010. ISSN 02646021. doi: 10.1042/BJ20100862.

44. G. Ekin Atilla-Gokcumen, Eleonora Muro, Josep Relat-Goberna, Sofia Sasse, Anne Bedigian, Margaret L. Coughlin, Sergi Garcia-Manyes, and Ulrike S. Eggert. Dividing cells regulate their lipid composition and localization. Cell, 156(3):428–439, 2014. ISSN 00928674. doi: 10.1016/j.cell.2013.12.015.

45. Schuyler B. Van Engelenburg, Gleb Shtengel, Prabuddha Sengupta, Kayoko Waki, Michal Jarnik, Sherimay D Ablan, Eric O Freed, Harald F Hess, and Jennifer Lippincott-Schwartz. Distribution of ESCRT machinery at HIV assembly sites reveals virus scaffolding of ES-CRT subunits. Science, 343(6171):653–656, 2014. ISSN 10959203. doi: 10.1126/science.1247786.

46. Jane F. Kaye and Andrew M. L. Lever. trans-Acting proteins involved in RNA encapsidation and viral assembly in human immunodeficiency virus type 1. Journal of virology, 70(2): 880–886, 1996. ISSN 0022-538X. doi: 10.1128/jvi.70.2.880-886.1996.

47. Stephen Cobeldick. Natural-Order Filename Sort, 2019. MATLAB Central File Exchange, Retrieved September 30, 2018.

## Bibliography

1. Ignacio Arganda-Carreras, Verena Kaynig, Curtis Rueden, Kevin W Eliceiri, Johannes Schindelin, Albert Cardona, and H Sebastian Seung. Trainable Weka Segmentation : a machine learning tool for microscopy pixel classification. Bioinformatics, 33(March):2424–2426, 2017. doi: 10.1093/bioinformatics/btx180.

